# CLEAR: Concise List Enrichment Analysis Reducing Redundancy

**DOI:** 10.64898/2026.03.30.715378

**Authors:** Xinglin Jia, An Phan, Karin Dorman, Claus Kadelka

## Abstract

**Motivation:** High-throughput experiments generate genome-wide measurements for thousands of genes, which are often tested marginally. Biological processes are driven by coordinated groups of genes rather than individual genes, making gene set enrichment analysis an essential *post hoc* interpretation tool. Traditional approaches such as Over-Representation Analysis and Gene Set Enrichment Analysis test gene sets independently, which ignores the hierarchical and overlapping structure of gene set collections such as the Gene Ontology, and often leads to redundant enrichment results. Set-based approaches such as MGSA address this issue by modeling multiple gene sets simultaneously, but they rely on binary gene activation states derived from arbitrary thresholds on gene-level statistics.

**Results:** We introduce Concise List Enrichment Analysis Reducing Redundancy (CLEAR), a Bayesian gene set enrichment framework that jointly models gene sets while incorporating continuous gene-level statistics such as test statistics or p-values. CLEAR extends model-based gene set analysis by replacing threshold-based gene activation with a probabilistic model for continuous gene-level statistics. This approach preserves the redundancy-reduction advantages of set-based enrichment methods while avoiding the information loss introduced by binarization. Using both simulated datasets and human gene expression data, we show that CLEAR improves sensitivity compared with existing enrichment approaches while producing a more concise and interpretable set of enriched gene sets.

**Availability and implementation:** The source code, data, and a brief tutorial are freely available at https://github.com/jiatuya/CLEAR

## 1 Introduction

Many modern high-throughput experiments generate genome-, transcriptome-, or proteome-wide measurements for thousands of genes. Hypotheses are often assessed marginally, but biological functions are typically coordinated by groups of genes rather than individual genes. Consequently, gene set analysis, also known as functional enrichment analysis, has become a fundamental tool for interpreting system-wide molecular data.

Early enrichment methods such as Over-Representation Analysis (ORA) test whether individual gene sets contain more genes of interest (e.g., differentially expressed genes) than expected by chance using statistical tests such as the hypergeometric or Fisher’s exact test [Khatri et al., 2012, Maleki et al., 2020]. ORA remains widely used and is often the primary option in several popular bioinformatics tools, including DAVID [Huang et al., 2009], WebGestalt [Zhang et al., 2005], and Enrichr [Chen et al., 2013, Kuleshov et al., 2016]. Gene Set Enrichment Analysis (GSEA) introduced a ranking-based framework that avoids the need to define a threshold for genes of interest by testing whether genes in a set tend to cluster at the extreme of a genome-wide ranked list [Subramanian et al., 2005]. Because GSEA utilizes continuous gene-level statistics such as fold changes or test statistics, it has become one of the most widely adopted enrichment methods for analyzing genomics, transcriptomics, and proteomics data.

Despite their widespread use, ORA and GSEA analyze each gene set independently. This assumption is problematic for gene set collections with hierarchical and overlapping relationships, such as the Gene Ontology (GO) [Tarca et al., 2013, Ashburner et al., 2000, Consortium, 2025]. In such cases, closely related parent and child gene sets often appear simultaneously in enrichment results, producing highly redundant lists that are difficult to interpret. To address this limitation, set-based approaches such as GenGO [Lu et al., 2008] and Model-based Gene Set Analysis (MGSA) [Bauer et al., 2010] jointly model the activation of multiple gene sets using probabilistic frameworks. By accounting for gene set overlap, these methods aim to identify a concise set of active biological processes rather than long lists of redundant gene sets.

However, existing set-based methods rely on binary gene activation states derived from arbitrary threshold on gene-level statistics. This binarization discards substantial information contained in the continuous statistics produced by upstream differential expression analyses, such as effect sizes or *p*-values. Some extensions of MGSA incorporate the topological structure of the Gene Ontology graph [Frost and McCray, 2012, Sun et al., 2017] or additional constraints [Wang et al., 2015], yet they still disregard the relevant continuous statistical information associated with the genes.

Here we introduce Concise List Enrichment Analysis Reducing Redundancy (CLEAR), a Bayesian framework that directly models the continuous gene-level statistics rather than binary activation states. CLEAR assumes that gene-level statistics follow distinct distributions under the null and alternative hypotheses and jointly infers the activation of gene sets while accounting for overlapping annotations. By leveraging the full statistical information available for each gene, CLEAR provides a more sensitive and nuanced characterization of gene set activity while maintaining the ability to reduce redundancy among related gene sets.

We evaluate CLEAR using both simulated data and real human gene expression datasets, and compare its performance to established enrichment methods, including ORA, GSEA, and MGSA. Across a range of scenarios, CLEAR improves sensitivity while preserving the interpretability advantages of set-based models by returning concise collections of enriched gene sets. These results demonstrate that modeling continuous gene-level statistics within a joint probabilistic framework provides a powerful and flexible approach to gene set enrichment analysis.

## 2 Materials and Methods

### 2.1 Generative Model

Following the generative model described in MGSA and multi-functional analyzer (MFA) [Wang et al., 2015], we consider *n* genes and *m* predefined gene sets. The annotation structure, which determines the genes associated with each gene set, is represented by an incidence matrix *I* = (*I*_*ij*_), where *I*_*ij*_ = 1 if gene *i* belongs to gene set *j*, and *I*_*ij*_ = 0 otherwise. Each gene set *j* has an unobserved activation indicator *T*_*j*_ ∈ {0, 1}, where *T*_*j*_ = 1 if gene set *j* is active, for example if its master regulatory gene is active, *T*_*j*_ = 0 otherwise. We assume *T*_*j*_ follows a Bernoulli(*π*) distribution, where *π* controls the expected proportion of active gene sets.

As in MGSA and MFA, CLEAR assumes that the hidden state of a gene *i*, denoted as *H*_*i*_, is fully determined by the hidden state of gene sets ***T*** = (*T*_1_, *T*_2_, …, *T*_*m*_) and incidence matrix *I* = (*I*_*ij*_). That is, a gene is active if it belongs to at least one active gene set:

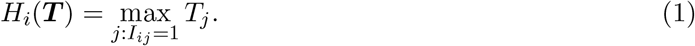

For each gene *i*, we observe a statistic *s*_*i*_ (e.g., a test statistic or *p*-value) that depends on its hidden state *H*_*i*_. Unlike MGSA and MFA, which model the binary state of *s*_*i*_, CLEAR models these observations through continuous null *f*_0_(*s* | *θ*_0_) and alternative distributions *f*_1_(*s* | *θ*_1_). That is,

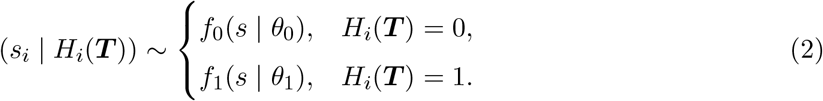

where *f*_0_ and *f*_1_ denote distribution families (e.g., Beta, Gamma, or Uniform) appropriate for the observed statistics (*p*-value or Wald test statistic), and *θ*_0_ and *θ*_1_ are the parameters of the distributions with priors

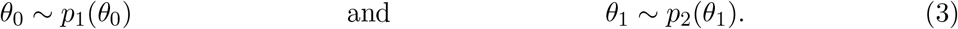

Therefore the likelihood of the observed statistics ***s*** = (*s*_1_, *s*_2_, …, *s*_*n*_) for all genes is:

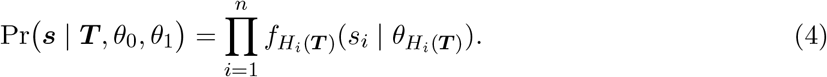

This generative formulation links gene-level statistics to latent gene set activity, allowing CLEAR to infer active biological processes directly from continuous gene-level measurements.

#### 2.1.1 Generative Model for Gene-level Statistics

Marginal differential expression (DE) analysis is a common upstream step in RNA-seq and other genomics studies, producing gene-level statistics such as test statistics or *p*-values. Methods such as DESeq2 assess differential expression using Wald tests on estimated log fold changes [Love et al., 2014]. Under the null hypothesis of no differential expression, the resulting Wald statistics approximately follow a standard normal distribution. Under the alternative hypothesis, the up- or down-regulated genes can be modeled as normally distributed with a shifted mean (*µ*_1_ ≠ 0) and increased variance 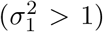 [Wolf et al., 2023]. Therefore, the combined distribution of up- and down-regulated genes is a mixture distribution with at least two components.

CLEAR models these gene-level statistics using distinct distributions under the null and alternative hypotheses. To limit the number of parameters it must estimate, CLEAR models the absolute value of Wald statistics using truncated Normal distributions, under both the null and the alternative (Table 1). This approach captures the increased magnitude of differential expression signals while avoiding separate modeling of up- and down-regulation. The same modeling assumption can also be applied to other DE analysis statistics, such as the moderated *t*-statistic from limma [Smyth, 2005].

**Table 1:** Distributions of *s*_*i*_ under null and alternative hypotheses (Wald statistics).

| Component | Description |
| --- | --- |
| $s_i$ | Absolute observed test statistic (Wald-like) |
| $f_0(s)$ | Truncated standard normal $\mathcal{N}_{[0,\infty)}(0, 1)$ |
| $f_1(s)$ | Truncated normal $\mathcal{N}_{[0,\infty)}(\mu_1, \sigma_1^2)$ |

When test statistics are unavailable or poorly approximated by Normal distributions, CLEAR can instead model *p*-values or transformed *p*-values (e.g., − log_10_(*p*)). Under the null hypothesis, *p*-values follow a uniform distribution on [0, 1], by definition. Under the alternative hypothesis, *p*-values concentrate near zero in a right-skewed distribution, which can be modeled by Beta distributions [Allison et al., 2002, Pounds and Morris, 2003].

In practice, transformed *p*-values may be easier to model. For instance, when sample sizes and effect sizes are sufficiently large, − log_10_(*p*) approximately follows a normal distribution under the alternative hypothesis [Boos and Stefanski, 2011], while under the null hypothesis it follows an exponential distribution with rate ln 10. Because − log_10_(*p*) is non-negative, the alternative distribution is modeled using a truncated Normal distribution defined on *s* ≥ 0.

In addition, CLEAR also offers an option to model − log(*p*)-values as Gamma-distributed under the alternative hypothesis, paired with Exp(1) under the null hypothesis. Compared to the truncated Normal, the Gamma distribution offers a natural generalization of the alternative model with flexible tail behavior, which is particularly useful when modeling asymmetric, right-skewed distributions.

These alternative likelihood models allow CLEAR to flexibly accommodate different forms of gene-level statistics while maintaining the same underlying probabilistic framework.

#### 2.1.2 Priors

Similar to MGSA, we placed a prior on the gene set activation probability *π* [Bauer and Gagneur, 2023]:

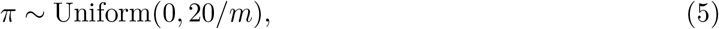

which reflects the prior belief that only a small fraction of gene sets are active a priori and encourages sparse gene set configurations. In addition, an empirical Bayes estimate of *π* can be obtained by maximizing the marginal likelihood; however, this quantity is difficult to compute. We show that the resulting estimate 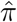 is close to the posterior mean proportion of active gene sets (Supplement A).

For *θ*_1_ we used weakly informative priors (Table 3) allowing wide parameter exploration while penalizing implausibly large values, following established practice in Bayesian modeling [Gelman, 2006, Gelman et al., 2008, Lemoine, 2019]. These priors accommodate the behaviors of the statistics under the alternative (e.g., *p*-values concentrated near zero).

For the [0, ∞)-truncated Normal alternative models for |*s*| or − log_10_(*p*), we assumed a Cauchy prior with empirically estimated location 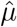 and scale of 2.5 for the location parameter *µ*,

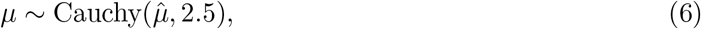

following Gelman and colleagues’ recommendation [Gelman et al., 2008, Bruch and Felderer, 2022]. And for *σ*, we used a half-Cauchy(0, 2.5) prior

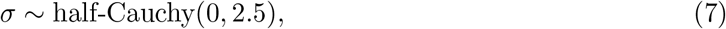

which is a widely recommended weakly informative prior for scale parameters to balance concentration near zero with heavy tails [Gelman, 2006, Polson and Scott, 2012, Bruch and Felderer, 2022].

We expect untransformed *p*-values under the alternative to be smaller than uniform and have a decreasing density on [0, 1] [Allison et al., 2002, Pounds and Morris, 2003, Gadbury et al., 2004, Gadbury and Allison, 2012]. Therefore, we choose a Beta(*a*, 1) distribution with 0 *< a <* 1, which is strictly decreasing on [0, 1]. We applied a noninformative Uniform prior on *a*:

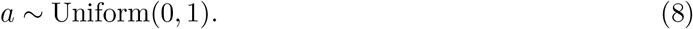

For the Gamma alternative for − log *p*-values, with shape *α* and scale *κ*, we use a weakly informative Exponential prior on *α*:

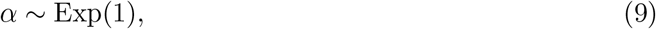

which creates a preference for shapes near the null (*α* = 1) while allowing flexibility. For the scale parameter *κ*, we again apply a half-Cauchy prior:

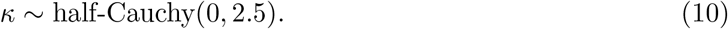

Under the null hypothesis, there are no free parameters *θ*_0_, and depending on user choice, *θ*_1_ is (*µ, σ*) of the truncated Normal, *a* of the Beta, or (*α, κ*) of the Gamma.

### 2.2 Markov Chain Monte Carlo (MCMC) estimation

We sample from the posterior distribution of gene set states ***T*** given the observed statistics ***s***,

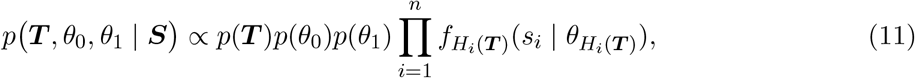

using a Metropolis-Hastings Markov Chain Monte Carlo (MCMC) algorithm [Robert and Casella, 2010]. As in MGSA, at each MCMC iteration, we update the gene set state with probability 0.8 and update the parameters of the alternative distributions with probability 0.2. Our MCMC algorithm is detailed here and illustrated in Fig. 1.

**Figure 1:**
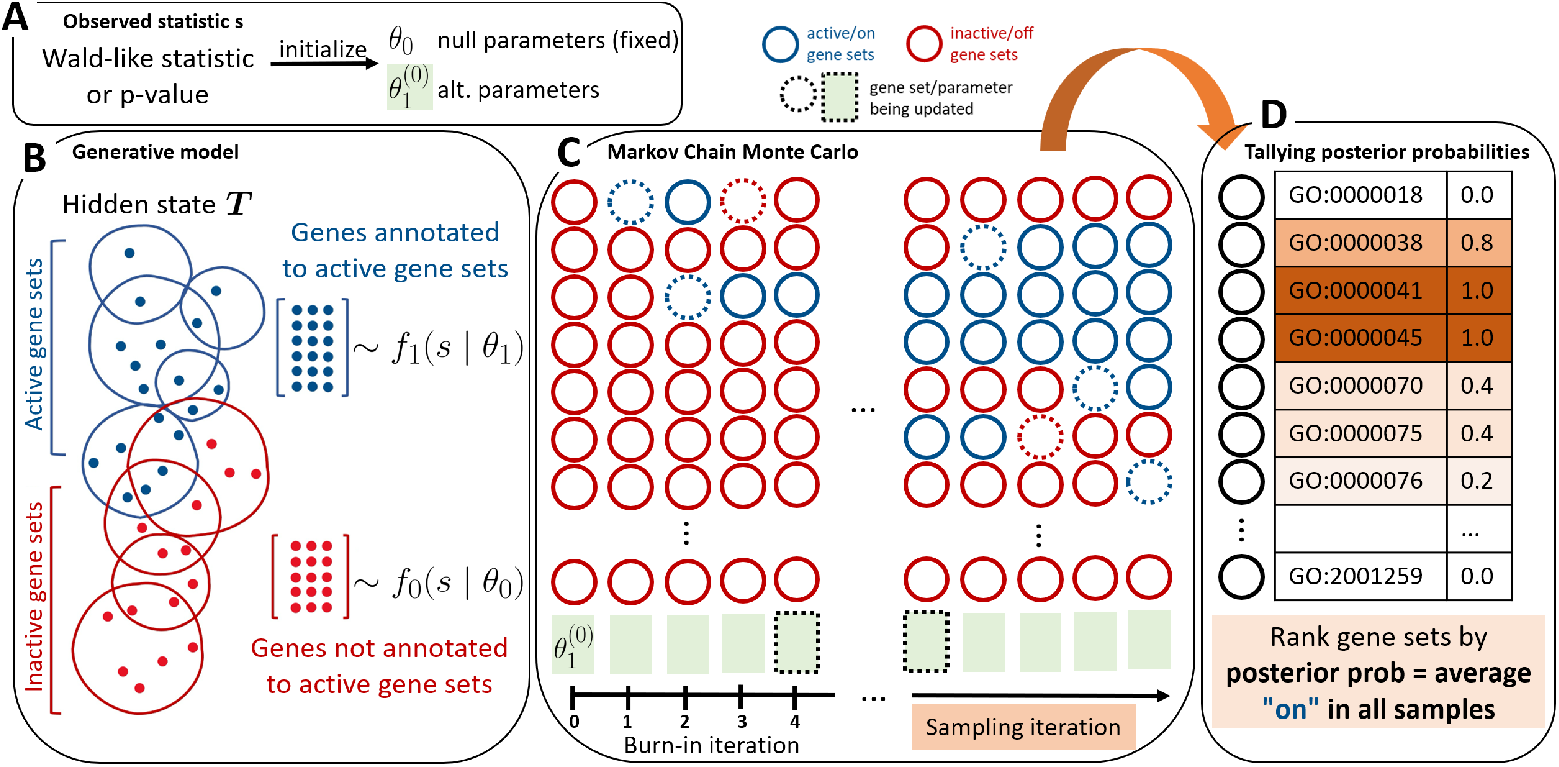
CLEAR method overview. CLEAR integrates a generative model for gene-level statistics with MCMC inference to identify active gene sets. (A) Observed gene-level statistics (Wald-like statistics or *p*-values) are used to initialize the parameters *θ*_0_ and 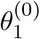. (B) Generative model of CLEAR. Each gene set *j* has an unobserved activation state *T*_*j*_; genes belonging to active gene sets follow the alternative distribution *f*_1_, while all other genes follow the null distribution *f*_0_. (C) MCMC sampling procedure. Starting from an initial configuration in which all gene sets are inactive, the algorithm iteratively updates the gene set states ***T*** and the distribution parameters *θ*_1_ (with probabilities 0.8 and 0.2, respectively) during burn-in and sampling. (D) Posterior probability of each gene set being active is computed as the proportion of sampling iterations in which the gene set is active/”on”.

#### 1. Initialize

An initial state ***T*** ^(0)^ is selected. In practice, we follow MGSA’s default and set ***T*** ^(0)^ ≡ 0, meaning all gene sets are initially inactive. We initialize the distribution parameters *θ*_1_ using simple summary statistics computed from the full set of observed statistics, which provides a neutral starting point that lies between the null and alternatives and avoids implausibly extreme values (Fig. 1A, Table 2).

**Table 2:** Initialization and update rules for alternative distributions.

| Type | Truncated Normal (\|Wald statistics\| or $-\log_{10}(p)$) | Beta ($p$) | Gamma ($-\log p$) |
| --- | --- | --- | --- |
| Deterministic | $\mu^{(0)} = \hat{\mu}, \sigma^{(0)} = \hat{\sigma}$ | $a^{(0)} = 0.5, b^{(0)} = 1$ | $\alpha^{(0)} = \frac{\hat{\mu}^2}{\hat{\sigma}^2}, \kappa^{(0)} = \frac{\hat{\sigma}^2}{\hat{\mu}}$ |
| Random | $\theta_1^{(0)} = \theta_1 u, \quad u \sim \text{Uniform}(0.5, 1.5)$ | | |
| Update rule | Propose one parameter at a time: $\theta'_{1j} \sim N(\theta_{1j}^{(k)}, \tau_j^2)$; re-propose if boundaries are violated | | |

*Note.* The quantities $\hat{\mu}$ and $\hat{\sigma}$ are the empirical mean and standard deviation of the transformed statistics appropriate to each model (e.g., |Wald statistics|, $-\log_{10}(p)$ , or $-\log(p)$ ). For the Gamma alternative, $(\alpha, \kappa)$ denote the shape and scale, and their initial values are obtained using method-of-moments estimates. The scalar $u$ is sampled from $\text{Uniform}(0.5, 1.5)$ and applied to $\theta_1$ for random initialization.

**Table 3:**
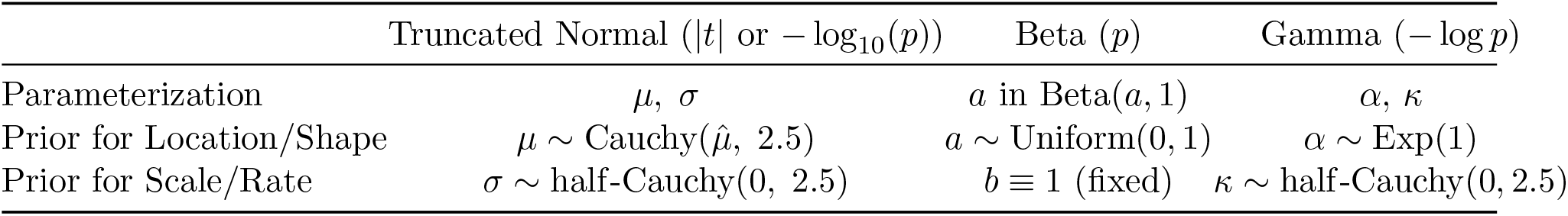
Prior distributions for alternative model parameters.

| | Truncated Normal ($\vert t\vert $ or $-\log_{10}(p)$) | Beta ($p$) | Gamma ($-\log p$) |
| --- | --- | --- | --- |
| Parameterization | $\mu, \sigma$ | $a$ in Beta($a, 1$) | $\alpha, \kappa$ |
| Prior for Location/Shape | $\mu \sim \text{Cauchy}(\hat{\mu}, 2.5)$ | $a \sim \text{Uniform}(0, 1)$ | $\alpha \sim \text{Exp}(1)$ |
| Prior for Scale/Rate | $\sigma \sim \text{half-Cauchy}(0, 2.5)$ | $b \equiv 1$ (fixed) | $\kappa \sim \text{half-Cauchy}(0, 2.5)$ |

#### 2. Updates of *T*

At each iteration *k*, the current gene set state is denoted as ***T*** ^(*k*)^. If iteration *k* is a gene set update, we propose a new state ***T*** ^*′*^ with proposal probability

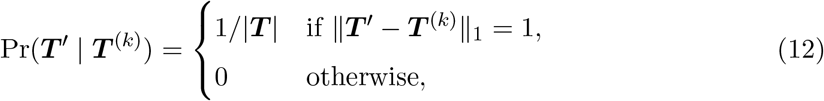

where |***T*** | is the number of gene sets. That is, we propose to flip the state of a single gene set by either activating or deactivating it.

#### 3. Updates of *θ*_1_

If iteration *k* is a distribution parameter update, we use a Gaussian random-walk proposals centered at the current values:

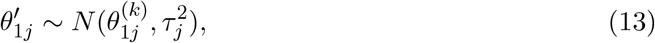

where *j* = 1, 2 indexes the elements of *θ*_1_ and *τ*_*j*_ is a proposal standard deviation tuned adaptively during burn-in based on proposal acceptance ratio to improve mixing [Andrieu and Thoms, 2008, Maire et al., 2019, Roberts and Rosenthal, 2009]. Every 1000 iterations during the burn-in period, if the acceptance ratio exceeds 0.45, *τ* is multiplied by 1.1; if it falls below 0.25, it is multiplied by 0.9. The specific thresholds and multipliers are arbitrarily chosen, but fall in the general recommended ranges [Roberts and Rosenthal, 2001]. Parameter proposals that violate the parameter constraints are rejected (e.g., *σ* ≤ 0, *α* ≤ 0, *κ* ≤ 0, or *a* ∉ (0, 1)).

This sampling scheme jointly infers gene set activity and the distribution of gene-level statistics by alternating updates of the active gene sets and the parameters of the alternative distributions.

### 2.3 In silico data generation

We generated in silico data based on real Gene Ontology (GO) gene set annotation data. For each simulated dataset, we randomly selected *m*_on_ ∈ {5, 10, 20, 50} from 883 *E. coli* GO-annotated gene sets [Gene Ontology Consortium, 2024]. The selected gene sets were required to be non-hierarchically related so that no chosen gene set was a subset of another. All genes annotated to at least one selected gene set were labeled as “active”, while the remaining genes were labeled “inactive”.

Gene-level test statistics were generated by simulating *t*-statistics for the active and inactive genes. Active genes were sampled from a non-central t distribution *t*_*ν*_(*µ*_*t*_) with *µ*_*t*_ *>* 0, while inactive genes were sampled from *t*_*ν*_(0). The degrees of freedom *ν* were set to 50 and 3 to represent experiments with sufficient and limited sample sizes, respectively. The non-centrality parameter (NCP) *µ*_*t*_, which controls the effect size, was varied across the range *µ*_*t*_ ∈ {1, 2, 3, 5, 10} based on empirical distributions observed in human RNA-seq datasets (Fig. S6).

To mimic both up- and down-regulation, the sign of each simulated *t*-statistic was flipped with probability 0.5. Two-sided *p*-values were then computed from the simulated *t*-statistics. This simulation scheme mimics a typical differential expression analysis pipeline used in gene expression studies, where gene-wise *t*-statistics are computed prior to enrichment analysis. Because the effective sample size *ν* + 1 is typically constant across genes in differential expression tests, varying *µ*_*t*_ allows us to evaluate enrichment methods under different signal strengths.

### 2.4 Real data

#### 2.4.1 Datasets

We collected several datasets published in popular benchmarking papers, including 15 human RNA-seq datasets from The Cancer Genome Atlas [Geistlinger et al., 2020], and 24 human microarray datasets in Gene Expression Omnibus (GEO) [Tarca et al., 2013]. The microarray datasets were restricted to only those with phenotypically relevant gene sets determined in [Geistlinger et al., 2020]. A detailed list of all datasets used can be found in Table S1.

For the TCGA datasets, raw count data for each cancer dataset were downloaded using the `GSEABenchmarkeR` R package, version 1.24.0 [Geistlinger et al., 2023]. We retained only genes with a minimum of 10 counts in at least three samples, consistent with DESeq2 best practices [Love et al., 2014, 2025]. Differential expression analysis on 15 TCGA datasets was performed using `DESeq2` without *p*-value adjustment [Love et al., 2014].

For the GEO microarray datasets, we obtained already preprocessed and normalized expression values from the `KEGGdzPathwaysGEO` R package, version 1.42.0 [Bhatti and Tarca, 2024]. We performed differential expression analysis using `limma` following [Tarca et al., 2012, Smyth, 2005] to get moderated t-statistics and unadjusted *p*-values. The `limma` package computes moderated t-statistics with a shrunken standard error after estimating the variance with empirical Bayes [Ritchie et al., 2015]. After that, probes were mapped to gene symbols using either Affymetrix Affymetrix HG-U133A or HG-U133 Plus 2 Array annotation data in R, as recorded in the metadata of each microarray dataset [Carlson, 2021a,b].

We used the GO biological processes (obtained from GO Data Archive, 2024-11-03 release) [Carbon and Mungall, 2024]. GO annotations for all human genes were retrieved from the same time point. All annotations were propagated to the root following the true path rule [Ashburner et al., 2000, Consortium, 2025]. For each dataset, GO gene sets were intersected with measured genes (i.e., genes not present in the dataset were excluded from GO gene sets for that dataset), and equivalent GO gene sets were combined into a single GO gene set (e.g., `mitotic spindle elongation` and `mitotic spindle midzone assembly` were combined into `mitotic spindle elongation & mitotic spindle midzone assembly` since they annotate the same genes). After all preprocessing steps, we retained approximately 3,000 GO gene sets of sizes between 20 and 500 genes. The upper gene set size cutoff removes overly broad and generic processes, such as `reproduction`, that do not pinpoint meaningful biological signals in the data. Lower gene set size cutoff eliminates very specific processes that are associated only with a few genes because they may represent noise rather than biological signals. Note that most processes have only very few gene annotations so the use of a lower cutoff dramatically decreases the number of considered gene sets and improves run time.

MGSA and GSEA were performed in R with Bioconductor packages (`mgsa` version 1.50 and `clusterProfiler::GSEA` version 4.10) [Bauer and Gagneur, 2023, Wu et al., 2021, 2023]. Scripts to run hypergeometric method and CLEAR are available on our GitHub repository. For MGSA and CLEAR, all datasets were run with 1,000,000 iterations (the first 500,000 are burn-in iterations), gene set states and hyperparameters are updated with probabilities 0.8 and 0.2, respectively. For all datasets, unadjusted *p*-values were used with the following methods: CLEAR tnormal (truncated normal), CLEAR beta, CLEAR gamma, MGSA, hypergeom (ORA). Wald statistics (from RNA-seq datasets) or moderated *t*-statistics (from microarray datasets) were used in GSEA and CLEAR tnormal.

All seven methods were provided with the same list of genes and GO gene sets to enable a fair comparison of the enrichment analysis output. For all variants of CLEAR, we repeated the analysis ten times for each dataset (to reduce variability from MCMC initialization), and ranked the final list of gene sets by highest sum of posterior probability over all repeats.

#### 2.4.2 Metrics

To assess biological relevance in real datasets, we evaluated how well each method enriched known disease-relevant biological processes. We obtained a biological “ground truth” based on known disease driver genes and oncogenic processes [Geistlinger et al., 2020]. Disease-relevant gene sets were extracted from the MalaCards disease database, which measures disease relevance of genes from experimental evidence and the literature [Rappaport et al., 2014]. In our study, cancer- and disease-relevant gene sets with 20 to 500 associated genes were considered positives, while other gene sets of the same size range were treated as negatives. The number of disease-relevant gene sets for each dataset is reported in Table S1.

Gene set–wise precision-recall curves were constructed using the positive and negative gene sets, and performance was summarized by the area under the precision–recall curve (PR-AUC). Following standard practice, the curve was anchored by projecting the precision at the first non-zero recall to recall = 0 and extending to recall = 1 with precision equal to the prevalence (i.e., the fraction of positive gene sets). Gene sets returned by each method were ranked either by highest posterior probability (for MGSA and CLEAR), or by lowest *p*-value (for GSEA and ORA). Additionally, since each dataset has a different number of phenotypically relevant gene sets (ranging from 6 to 282 gene sets, see Table S1), we normalized the PR-AUC by the prevalence of each dataset. Under this normalization, a random ranking of gene sets yields a normalized PR-AUC of 1.0.

Since we compare ORA and GSEA with set-based methods (MGSA and CLEAR), we also assessed the overlap of the top-ranking gene sets to emphasize the advantage of analyzing all gene sets simultaneously. Set-based methods are expected to select gene sets that share fewer genes, avoiding the long lists of overlapping gene sets commonly produced by independent testing approaches [Jantzen et al., 2011, Simillion et al., 2017]. To quantify this property, we selected the top 20 highest-ranked gene sets from each method (breaking ties randomly when necessary), and computed the pairwise overlap coefficient between gene sets based on their annotated genes [Candia and Ferrucci, 2024]. The average of all pairwise overlap coefficients defines the *gene set overlap* metric used in this study. In addition, we also calculated the average size of the top 20 gene sets returned by each method, which serves as a proxy for the specificity of the enriched biological processes.

Performance differences across enrichment methods were assessed using the Friedman test, treating datasets as blocks and methods as repeated/grouped measures. This non-parametric test accounts for grouped measurements across datasets and does not assume normality of the performance metric [Friedman, 1937]. When the global test was significant, we applied the Nemenyi test for post-hoc pairwise comparisons [Nemenyi, 1963].

## 3 Result

We evaluated the performance of CLEAR against MGSA, ORA, and GSEA on both in silico and real data [Bauer and Gagneur, 2023, Subramanian et al., 2005, Zhang et al., 2005, Huang et al., 2009]. The data contain gene sets ranked either by descending posterior probability (CLEAR and MGSA) or increasing *p*-value (ORA and GSEA). For in silico datasets, precision-recall (PR) curves and redundancy were used to capture the power and accuracy of the methods.

### 3.1 In silico data analysis

PR curves measure how effectively a method recovers true positive gene sets. Precision is the proportion of true positives among all returned positives, while recall (also known as sensitivity) is the proportion of true positives identified out of all positives. For each simulated dataset, we computed the area under the PR curve (PR-AUC) to summarize overall performance.

Across simulated datasets, CLEAR consistently achieved higher PR-AUC than existing enrichment methods, particularly under moderate to strong signal conditions. The improvement was most pronounced when gene-level statistics closely followed the assumed likelihood distributions (Fig. 2).

**Figure 2:**
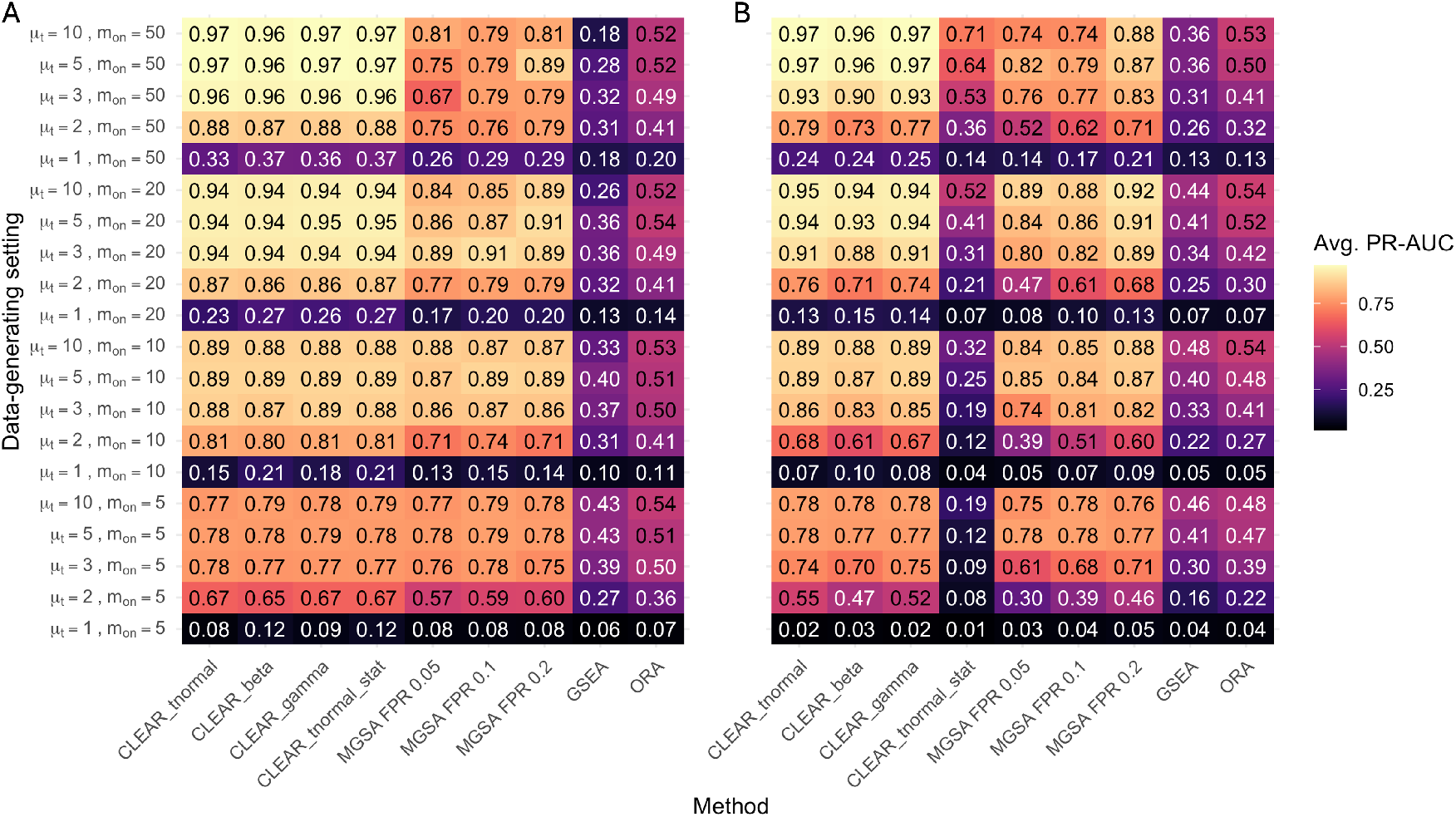
Average precision-recall AUC (PR-AUC) of enrichment methods in simulated data. Rows describe different data-generation configurations, characterized by the signal strength *µ*_*t*_ and the number of activated gene sets *m*_on_. Columns describe enrichment methods. Panel A shows results for a sufficient sample size (*ν* = 50). Panel B shows results for a limited sample size (*ν* = 3).

When the sample size is sufficiently large (*ν* = 50), all variants of CLEAR (truncated Normal (*p*-values), Beta, Gamma, and truncated Normal (test statistics)) performed equally well with highest PR-AUC among all methods being tested, followed by *p*-value-based models, MGSA, ORA, and then GSEA (Fig. 2A). Compared to MGSA and the traditional methods, CLEAR showed a clear advantage when the signal-to-noise ratio was high (e.g., *µ*_*t*_ = 10). When the signal is very weak (e.g., *µ*_*t*_ = 1), all methods perform poorly and the performance gain of *p*-value-based CLEAR models over MGSA was smaller.

When the sample size is small (*ν* = 3), performance declines for all methods due to increased statistical noise. The test statistics model was particularly affected because the Normality assumption for the alternative distribution no longer holds well in small-sample settings. In contrast, the *p*-value-based CLEAR models maintained strong performance and consistently outperformed other methods in most scenarios.

Overall, CLEAR variants substantially outperformed MGSA, ORA, and GSEA when the signal strength was moderate to high. Under very weak signals (e.g., NCP = *µ*_*t*_ = 1), all methods performed poorly, and the differences between methods were smaller.

### 3.2 Real data analysis

We first examined the characteristics of gene sets returned by each enrichment method. CLEAR and GSEA tended to return larger gene sets than MGSA and ORA (Fig. 3A). For CLEAR, this behavior likely reflects the hierarchical structure of the GO graph. When both a parent gene set and its child gene sets carry detectable signal, CLEAR often selects the parent gene set while excluding the children.

**Figure 3:**
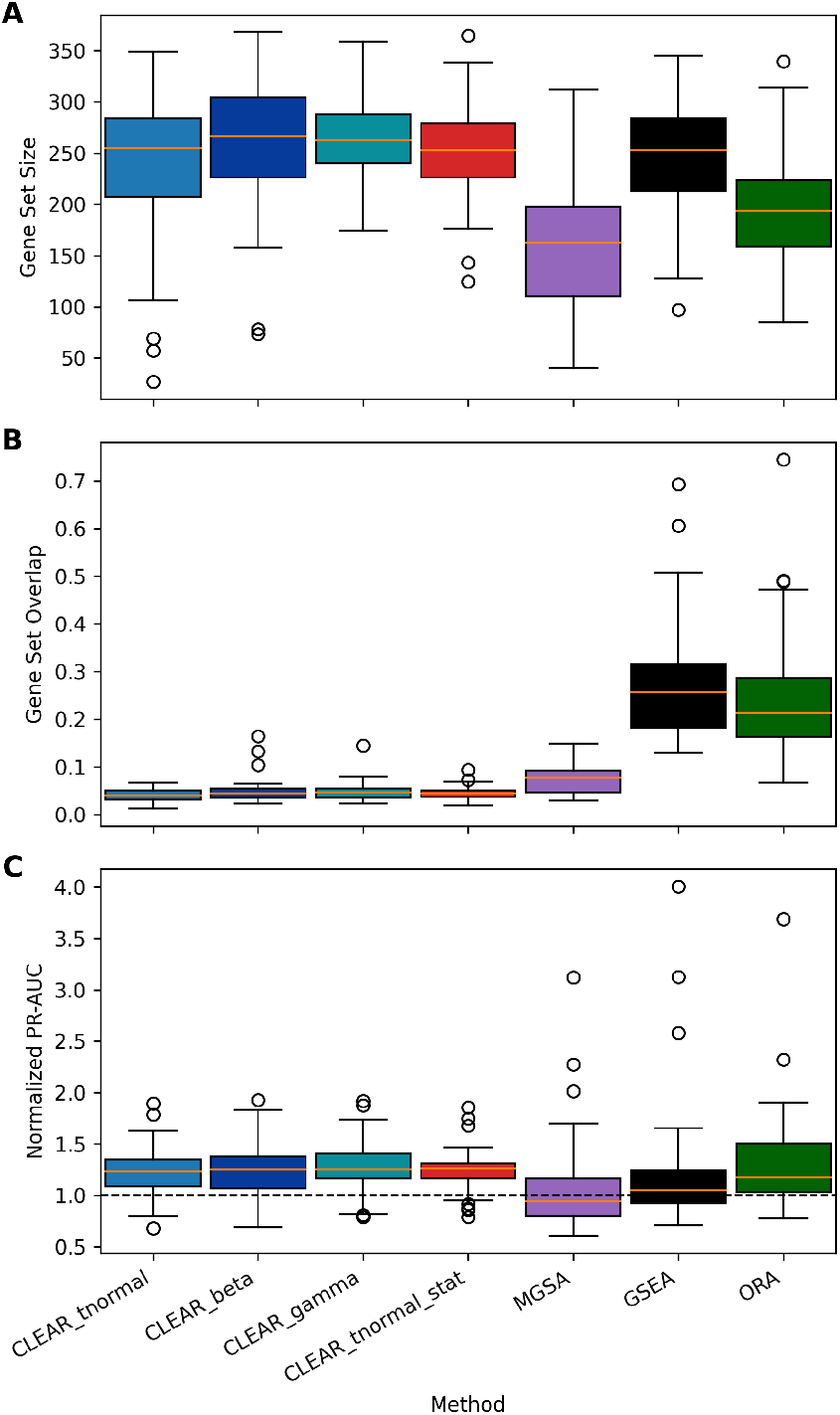
Method performance across 15 TCGA RNA-seq datasets and 24 GEO microarray datasets. Panels A and B show the average gene set size and gene set overlap for the top 20 gene sets enriched by each method. Panel C shows the normalized PR-AUC values where phenotypically relevant gene sets are considered positives. Orange lines display the median value for each method. Dotted black line in panel C represents the expected value of normalized PR-AUC.

Next, we evaluated redundancy among the top enriched gene sets (Fig. 3B). CLEAR produced the lowest gene set overlap, followed by the other set-based method MGSA. In contrast, the traditional enrichment methods GSEA and ORA returned substantially more overlapping gene sets. Notably, CLEAR achieved the lowest overlap despite tending to return relatively large gene sets (Fig. 3A), indicating that CLEAR effectively selects representative gene sets rather than multiple closely related gene sets.

Finally, we assessed biological relevance using normalized PR-AUC scores, which measure how effectively each method recovers known disease-relevant biological processes (Fig. 3C). Normalization allows comparison across datasets with different proportions of positive gene sets. Raw PR-AUC values are reported in the Supplement (Fig. S5). For most datasets, all methods performed only slightly better than a random prediction, hinting at very low signal-to-noise ratios. The Friedman global test [Friedman, 1937], using datasets as the grouping variable, rejected the null hypothesis that all methods performed similarly (*p* = 1.6 × 10^−10^). CLEAR achieved performance comparable to ORA (*p* ≈ 1.0, Nemenyi post-hoc test), implying that CLEAR performed no worse than ORA [Woolson, 2007, Benavoli et al., 2016]. All versions of CLEAR significantly outperformed MGSA and GSEA (*p <* 0.03, Nemenyi post-hoc test), except for CLEAR tnormal versus GSEA (*p* = 0.26). All post-hoc test results are reported in Fig. S7.

It is important to note that the disease-relevant biological processes used as positives in the normalized PR-AUC metric are often hierarchically related. For example, in the BRCA dataset the gene set `cell surface receptor protein tyrosine kinase signaling pathway (GO:0007169)` and three of its child gene sets `GO:0008543, GO:0038084, GO:0048009` are all considered positives. CLEAR selected only the parent gene set, whereas GSEA and ORA selected all four gene sets, thereby receiving four true positives in the precision-recall metric. This evaluation framework therefore favors methods that return redundant gene sets. Despite this disadvantage, CLEAR achieved comparable PR-AUC to ORA and better PR-AUC than MGSA, and GSEA, and a hypothetical random prediction. Overall, CLEAR consistently returned less redundant gene sets while achieving comparable or better biological relevance than traditional enrichment methods.

We evaluated elapsed runtime and memory usage for all methods on two human RNA-seq datasets, BLCA (3,304 GO gene sets) and BRCA (3,334 GO gene sets), using the GNU time utility running on a single CPU. Among the evaluated methods, CLEAR had the longest runtime. For one run of 1,000,000 iterations, CLEAR beta and CLEAR gamma required approximately 20 minutes, whereas CLEAR tnormal (using either *p*-value or Wald statistic input) required approximately 10 minutes. Under the same MCMC iteration setting, MGSA completed in a few seconds with an efficient implementation in C. GSEA with the fgsea preranked implementation required approximately 2 minutes, and ORA completed in a few seconds per dataset [Korotkevich et al., 2021]. Although CLEAR is computationally slower than methods such as ORA, GSEA, and MGSA, this is not unexpected for a Bayesian-based approach that models all gene sets simultaneously and updates a high-dimensional joint state space through MCMC.

## 4 Discussion

We developed a novel enrichment analysis method, CLEAR, that extends set-based gene set analysis to directly model continuous gene-level statistics. By replacing binary gene activation with a probabilistic model for gene-level test statistics or *p*-values, CLEAR retains the redundancy-reduction advantages of set-based enrichment approaches while avoiding the information loss introduced by thresholding.

Across simulated datasets, CLEAR substantially outperformed existing enrichment methods under moderate to strong signal conditions. In real biological datasets (with seemingly much lower signal-to-noise ratios), CLEAR achieved equal and better biological relevance than traditional enrichment methods (GSEA, ORA, MGSA) while returning substantially fewer redundant sets of enriched GO gene sets. Together, these results demonstrate that modeling gene-level statistics directly can improve sensitivity while maintaining interpretable enrichment results.

In the simulation study, gene-level test statistics were generated from a *t*-distribution with different degrees of freedom *ν* to represent experiments with sufficient or limited sample sizes. This design mimics realistic differential expression analysis pipelines while avoiding biases that could arise from sampling statistics directly from assumed model distributions (e.g., mixtures of Beta and Uniform distributions for *p*-values). As a result, CLEAR does not gain an artificial advantage in the benchmarking experiments.

When the sample size is small (*ν* = 3), the simulated test statistics deviate from the truncated Normal assumption used in the statistic-based CLEAR model, leading to reduced performance (Fig. 2). In contrast, the *p*-value-based CLEAR models were more robust because the Uniform(0, 1) distribution assumed under the null model remained true regardless of the underlying test statistic distribution. Consequently, the *p*-value-based CLEAR models maintained strong performance even under small-sample conditions.

### CLEAR addresses MGSA’s binarization threshold issue

CLEAR extends MGSA’s binary generative framework (that relies heavily on the correct choice of arbitrary thresholds) by replacing threshold-based gene activation with a probabilistic model for threshold-free, continuous gene-level statistics. In MGSA, genes must be classified as active or inactive based on an arbitrary threshold, which can lead to information loss and sensitivity to threshold choice. CLEAR instead models gene-level statistics or *p*-values directly using parametric distributions under the null and alternative hypotheses. This approach avoids binarization and allows the model to capture more subtle gene-level signals.

CLEAR and MGSA belong to a broader class of set-based Bayesian enrichment approaches that jointly model gene sets rather than analyzing them independently, as done by traditional enrichment methods such as ORA and GSEA. By modeling gene sets jointly, set-based approaches address the redundancy and dependency inherent in gene set collections such as the Gene Ontology, reducing the tendency to return long lists of highly overlapping enriched gene sets.

### Benchmarking on real datasets remains challenging

CLEAR relies on differences between the gene-level statistics of genes under the alternative distribution (genes associated with the condition of interest) and those under the null distribution. In an ideal setting, genes belonging to phenotypically relevant gene sets would exhibit gene-level statistics clearly distinct from background genes. However, in the real datasets we observed substantial overlap between the *p*-value and Wald statistic distributions of genes belonging to phenotypically relevant gene sets and those outside them (Fig. S6). This pattern was observed across nearly all datasets, indicating that gene-level statistics alone often provide limited separation between benchmark-positive and benchmark-negative genes.

In addition, Fig. S6 also shows deviations from the model assumptions underlying CLEAR. In particular, the empirical distributions of benchmark-negative genes often differ from the assumed null distributions (Uniform for *p*-values in CLEAR beta, exp(1) for − log *p* in CLEAR gamma, and truncated Normal for |Wald statistics| in CLEAR tnormal). A likely explanation is that benchmark-negative genes in real data rarely represent a pure null set; instead, they may include genes with weak or indirect associations with the studied condition. In fact, many benchmark-negative genes exhibit unexpectedly small *p*-values, suggesting that curated phenotypically relevant gene sets are at best an imperfect proxy for true biological ground truth.

More broadly, benchmarking enrichment methods on real data is inherently challenging because definitive ground truth gene sets are rarely available. Instead, benchmarks rely on prior biological knowledge, which can be incomplete or inconsistent [Hung et al., 2012]. Even with carefully curated benchmarking datasets, it remains difficult to assess method performance in real biological systems such as cancer, where consensus pathway definitions are still evolving [Chen et al., 2021, Menyhart et al., 2024]. Similar challenges arise in simpler model organism experiments, where multiple biological processes may plausibly be considered relevant to the experimental condition [Siso et al., 2012, Bendjilali et al., 2017]. Our benchmarking therefore follows established practice in the literature while acknowledging that evaluations based on curated gene sets remain approximate in the absence of definitive biological ground truth.

### Limitations

Despite the advantages, CLEAR has a few limitations. First, MCMC convergence can be elusive given the high dimension of the parameter space. CLEAR jointly samples discrete and continuous parameters from a posterior distribution with potentially many local optima. MGSA also has the same issue, and its documentation recommends multiple restarts because convergence is difficult to access. We provide a detailed convergence analysis in Appendix B. Given this limitation, we recommend running multiple independent chains for each dataset to help confirm that the model has properly converged.

Second, runtime is a practical limitation of CLEAR. While ORA and GSEA evaluate each gene set individually, MGSA and CLEAR use a computationally intensive Bayesian framework that jointly models all gene sets. However, the current implementation of CLEAR takes longer than MGSA because it uses a more complex statistical framework. In MGSA, the parameters false positive rate and false negative rate lie in a bounded interval (0, 1), and MGSA defines a discrete grid over these parameters and pre-calculates the likelihood contributions for true positives, true negatives, false positives, and false negatives to be reused during MCMC and reduce computational cost. In contrast, CLEAR does not discretize parameters on a predefined grid because its continuous parameters (*θ*_1_ for *f*_1_) do not have similarly bounded ranges. As a result, the probability density of genes must be recomputed whenever the model parameters are updated during MCMC (20% of iterations), which becomes the main computational bottleneck. We also note that the current implementation of CLEAR is written in R, whereas MGSA is implemented in C, so part of the observed runtime difference likely arises from implementation efficiency rather than from the statistical framework alone. Future work should focus on improving computational efficiency through code profiling, lower-level implementation, and algorithmic optimization.

Third, CLEAR’s performance may be affected when empirical data deviate substantially from the assumed statistical distribution. CLEAR relies on predefined distributions to model the null and alternative hypotheses (e.g., Uniform and Beta). If the true distribution of the data differs from these assumptions, model fit accuracy may be reduced. In practice, different datasets may exhibit different mixture patterns in their gene-level statistics. Therefore, we recommend visually inspecting the distribution of gene-level statistics (e.g., *p*-values or test statistics) before fitting the model to achieve the best performance.

## 5 Conclusion

Existing functional enrichment methods often rely on independent testing of gene sets or binarized gene activation states, which can discard valuable information contained in continuous gene-level statistics. CLEAR addresses these limitations by modeling continuous gene-level statistics within a probabilistic framework that distinguishes between null and alternative gene distributions. Across simulated datasets, CLEAR demonstrated improved sensitivity compared with existing enrichment approaches. In real biological datasets, CLEAR achieved biological relevance comparable to traditional enrichment methods while producing substantially less redundant enrichment results. By combining the redundancy control of set-based enrichment approaches with a threshold-free probabilistic model for gene-level signals, CLEAR provides a practical, robust and interpretable framework for gene set enrichment analysis.

## Author contributions statement

Conceptualization: X.J., A.P., C.K. Methodology: X.J., A.P., K.D., C.K. Software: X.J., A.P. Formal analysis: X.J., A.P. Supervision: K.D., C.K. Writing – original draft: X.J., A.P. Writing – review & editing: X.J., A.P., K.D., C.K.

## Data availability

The source code and data are freely available at https://github.com/jiatuya/CLEAR. Publicly available datasets were used in this manuscript, as summarized in Section 2.4.1. Preprocessed datasets (TCGA RNA-seq and GEO microarray) are available in the Figshare folder at https://doi.org/10.6084/m9.figshare.31891432.

Supplementary methods and figures - Jia et al.

## A Empirical Bayes Interpretation of Gene set Activation Probability *π*

MGSA assumes that gene set activation indicators *T*_*j*_ ∈ {0, 1} follow

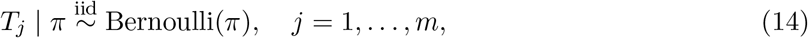

with a hyperprior distribution *π* ∼ Uniform(0, 20*/m*), where *m* is the number of gene sets. Let *s* denote the observed statistics and let *θ*_0_, *θ*_1_ denote the null and alternative parameters, respectively. In empirical Bayes, we can estimate *π* using marginal likelihood,

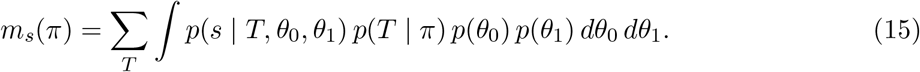

However, the marginal likelihood is generally difficult to compute. Instead, we consider the posterior distribution under a fixed value *π*,

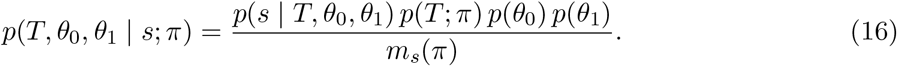

Suppose an MCMC sample 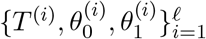 is obtained under a particular *π*_1_. Then the marginal likelihood ratio can be approximated by

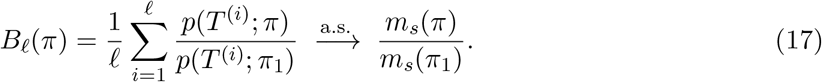

Therefore,

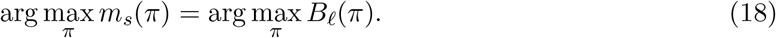

In the current setting, the maximum is

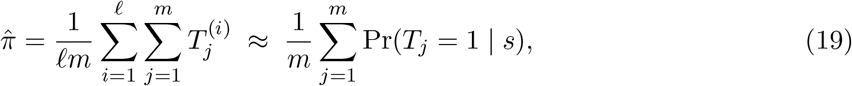

which shows that the empirical Bayes estimate of *π* approximates the posterior mean proportion of active gene sets.

## B Convergence Analysis

Convergence is an important indication that the posterior has been correctly estimated for models that rely on posterior sampling. For MGSA and CLEAR, the parameters include gene set activation, treated as a binary parameter, along with continuous parameters, such as false positive rate and false negative rate for MGSA, and the parameters *θ*_1_ of the alternative distributions for CLEAR. We employed separate strategies to evaluate the convergence of the continuous vs. discrete parameters. In addition, we used trace plots to monitor posterior changes over iterations.

### B.1 Discrete Parameter Convergence Assessment

We assessed the convergence of binary gene set indicators (0 for “off” and 1 for “on”) following Deonovic and Smith’s framework for categorical MCMC who offer two *χ*^2^ statistics for testing convergence of discrete parameters [Deonovic and Smith, 2017].

#### B.1.1 Homogeneity of the Marginal Probability using Weiss Test

We applied a Weiss test for convergence of posterior probabilities across all MCMC chains. The standard Pearson *χ*^2^ statistic was adjusted by a variance inflation factor, *ĉ*, to account for the dependence of MCMC sampling following the recommendations of Deonovic and Smith. Factor *ĉ* is estimated by modeling the binary sequence as a Discrete Autoregressive process of order 1 (DAR(1)).

#### B.1.2 Homogeneity of the Transition Probability using Billingsley Test

In addition to the Weiss test, we applied the Billingsley test to transition probabilities, i.e., 0 → 1, 1 → 1, etc., to assess convergence of chain dynamics. We constructed a contingency table of transition counts for each gene set and estimated chain-specific transition probabilities. Then we computed a *χ*^2^ statistic to compare these transition probabilities with the pooled transition probabilities across all chains.

The Weiss test and Bilingsley tests were performed for each gene set independently. We specifically focused the convergence analysis on “high-confidence” gene sets (gene sets with a posterior probability *>* 0.5) to ensure that the gene sets identified with highest support were robustly identified across independent runs.

### B.2 Continuous Parameter Convergence Assessment

For continuous parameters (e.g., mean and standard deviation from a truncated Normal), we assessed convergence using the standard 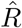 [Gelman and Rubin, 1992]. An 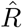 value below 1.1 was considered indicative of convergence.

### B.3 Convergence Result and Discussion

We performed convergence analysis on simulated datasets with *ν* = 50. We sampled 5 out of 100 replicates across all combinations of NCP and *m*_on_, totaling 100 datasets. For each dataset, we ran 5 independent MCMC chains and recorded the posterior probabilities of gene sets, transition probabilities, continuous parameter values, and the posterior value. The chi-square tests showed most high-confidence gene sets had non-significant *p*-values (*>* 0.05), indicating convergence of the binary sampling for those gene sets. However, in low-signal datasets, the proportion of converged gene sets tends to be lower than in high-signal datasets (Fig. S1). The 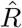 diagnostic showed good convergence of the continuous parameters for most datasets. However, lack of convergence was observed in the combinations of (NCP = 5, *m*_*on*_ = 5), (10, 5) and (10, 10) (Fig. S2).

**Figure S1:**
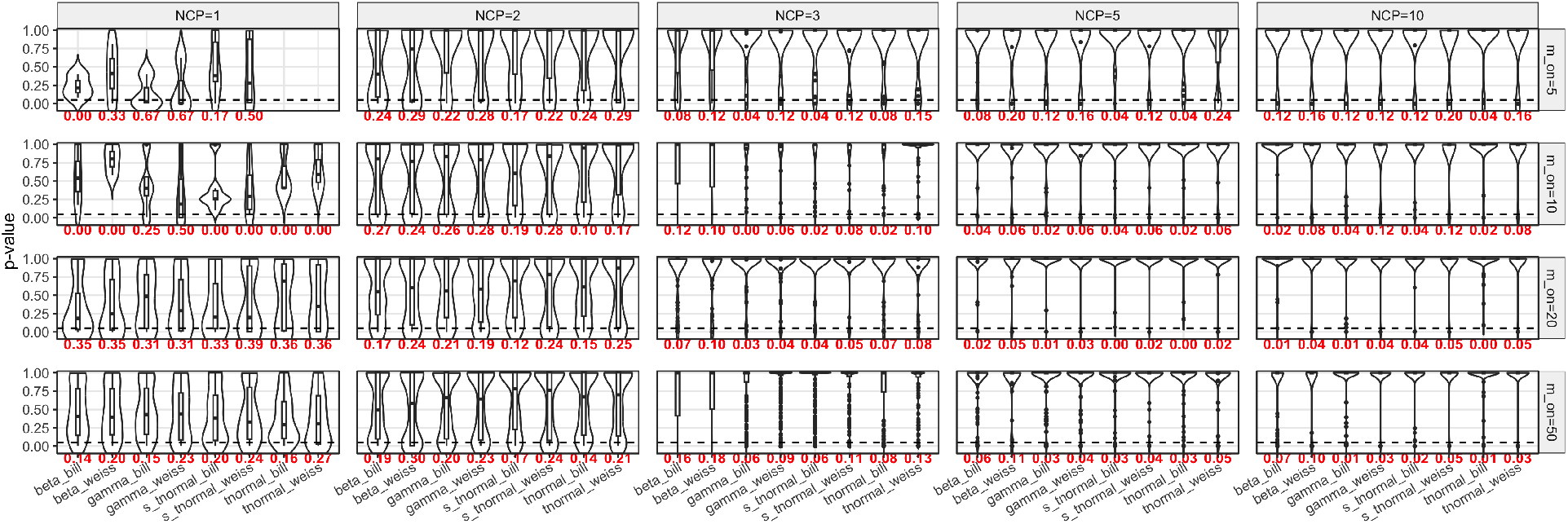
Weiss test and Billingsley test *p*-values for gene sets with average posterior probability *>* 0.5. Results are computed for simulated data sets under different data-generation configurations by the signal strength NCP (*µ*_*t*_) and the number of activated gene sets (*m*_on_). The dashed line marks the significance threshold (0.05). Red numbers at the bottom of each violin are the proportion of *p*-values below 0.05

CLEAR is a high-dimensional Bayesian model that samples from both discrete and continuous spaces. When the number of gene sets is high, it is challenging for CLEAR to fully explore the posterior distribution and converge to the global optimal, despite the large number of iterations. If multiple optimal or near-optimal solutions exist, different chains may converge to different regions of the posterior space. In the convergence analysis of the simulated datasets, we observed that some datasets having poor convergence metrics (i.e.,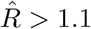) exhibited multimodal behavior, that different chains converged to different posterior configurations with one or more gene sets having noticeably different posterior probabilities across chains (Fig. S3, Fig. S4).

**Figure S2:**
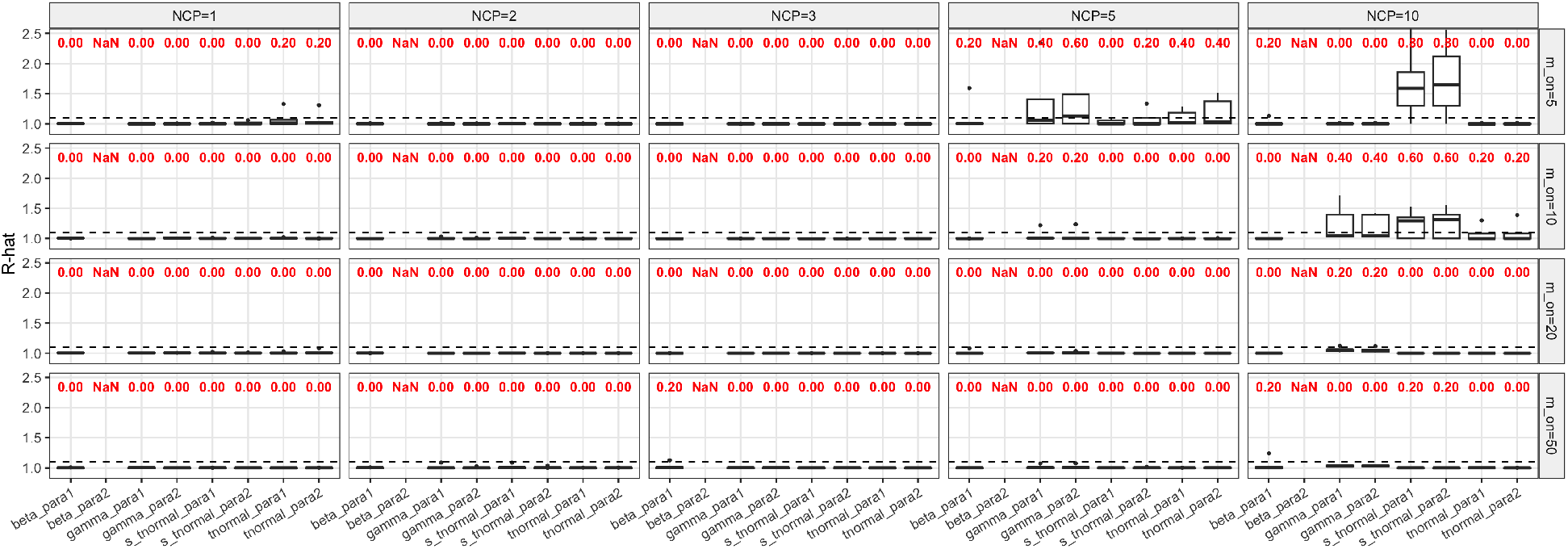
Gelman-Rubin convergence 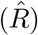 for each model parameter. Results are computed for simulated data sets under different data-generation configurations by the signal strength NCP (*µ*_*t*_) and the number of activated gene sets (*m*_on_). The dashed line marks the convergence threshold (1.1). Red numbers at the top of each violin are the proportion of 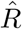 above 1.1

**Figure S3:**
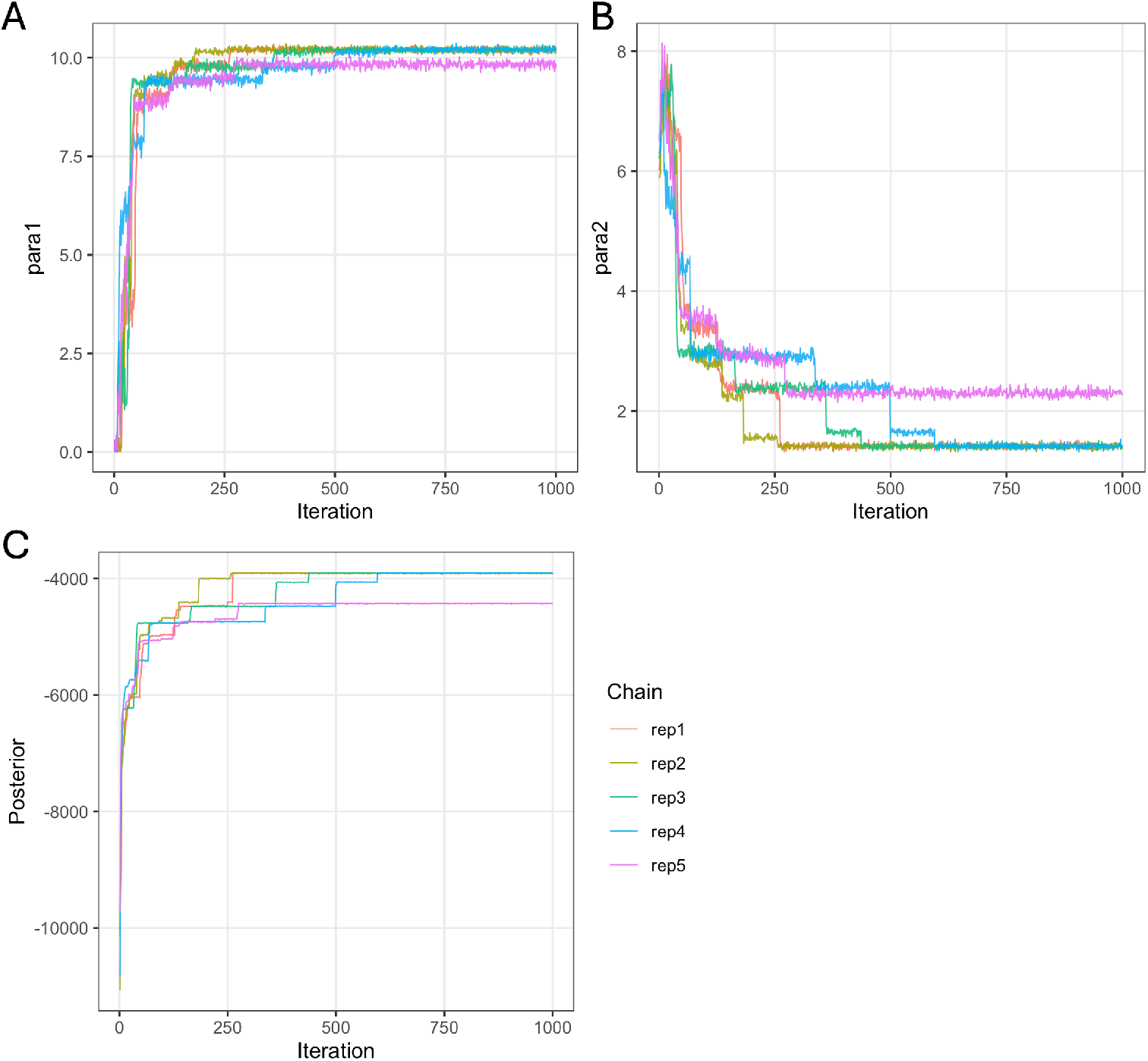
Convergence of CLEAR under the statistics model on a simulated dataset (*µ*_*t*_ = 10, *m*_on_ = 50). The trace plots record every 1000 MCMC steps from the beginning to the end. (A) Trace plot of *µ* across five MCMC chains. (B) Trace plot of *σ* across five MCMC chains. (C) Trace plot of posterior across five MCMC chains.

**Figure S4:**
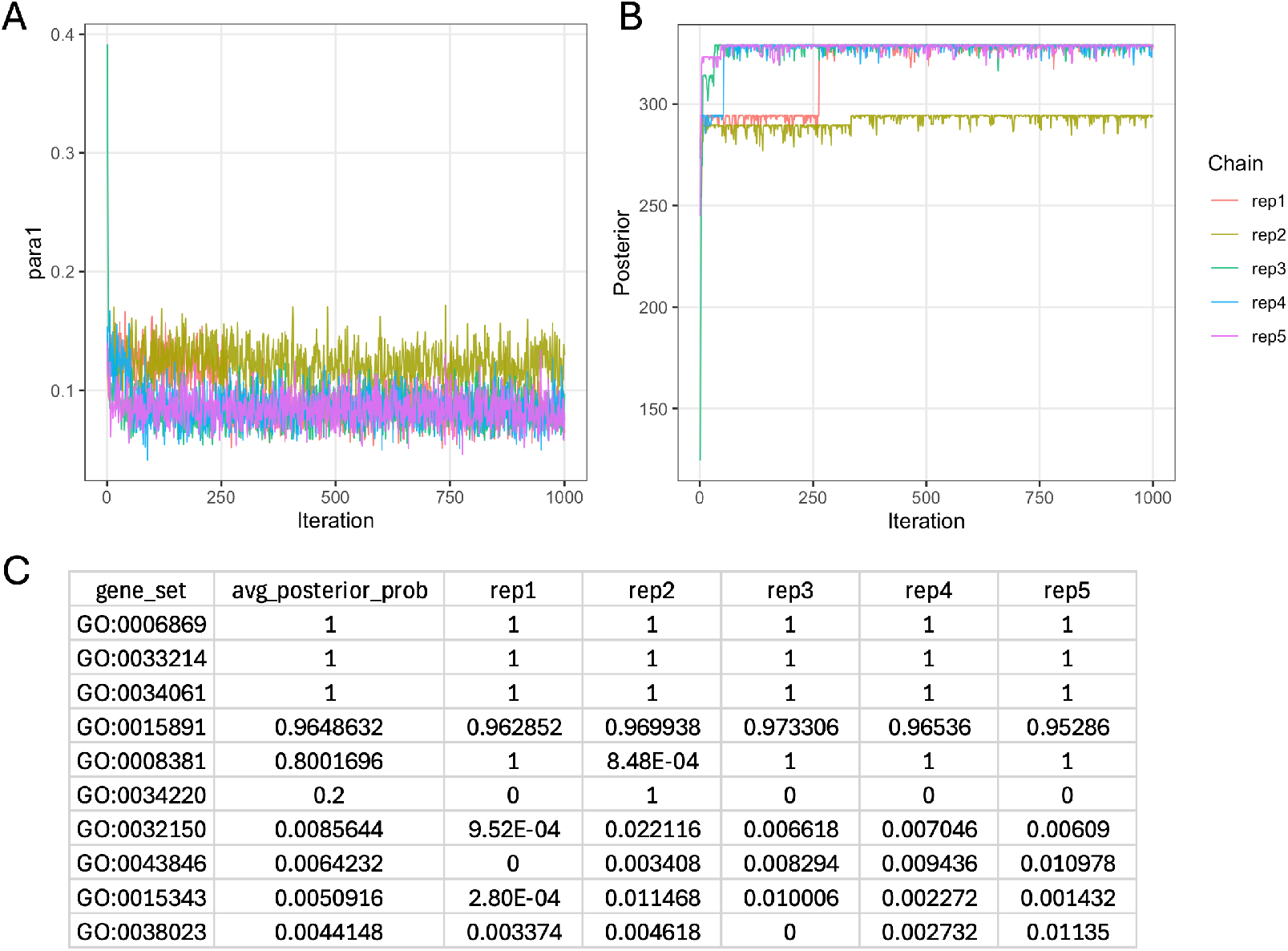
Convergence of CLEAR under the beta model on a simulated dataset (*µ*_*t*_ = 5, *m*_on_ = 5). The trace plots record every 1000 MCMC steps from the beginning to the end. (A) Trace plot of *a* across five MCMC chains. (B) Trace plot of posterior across five MCMC chains. (C) Posterior probabilities of the top gene sets ranked by average posterior probability across chains.

**Figure S5:**
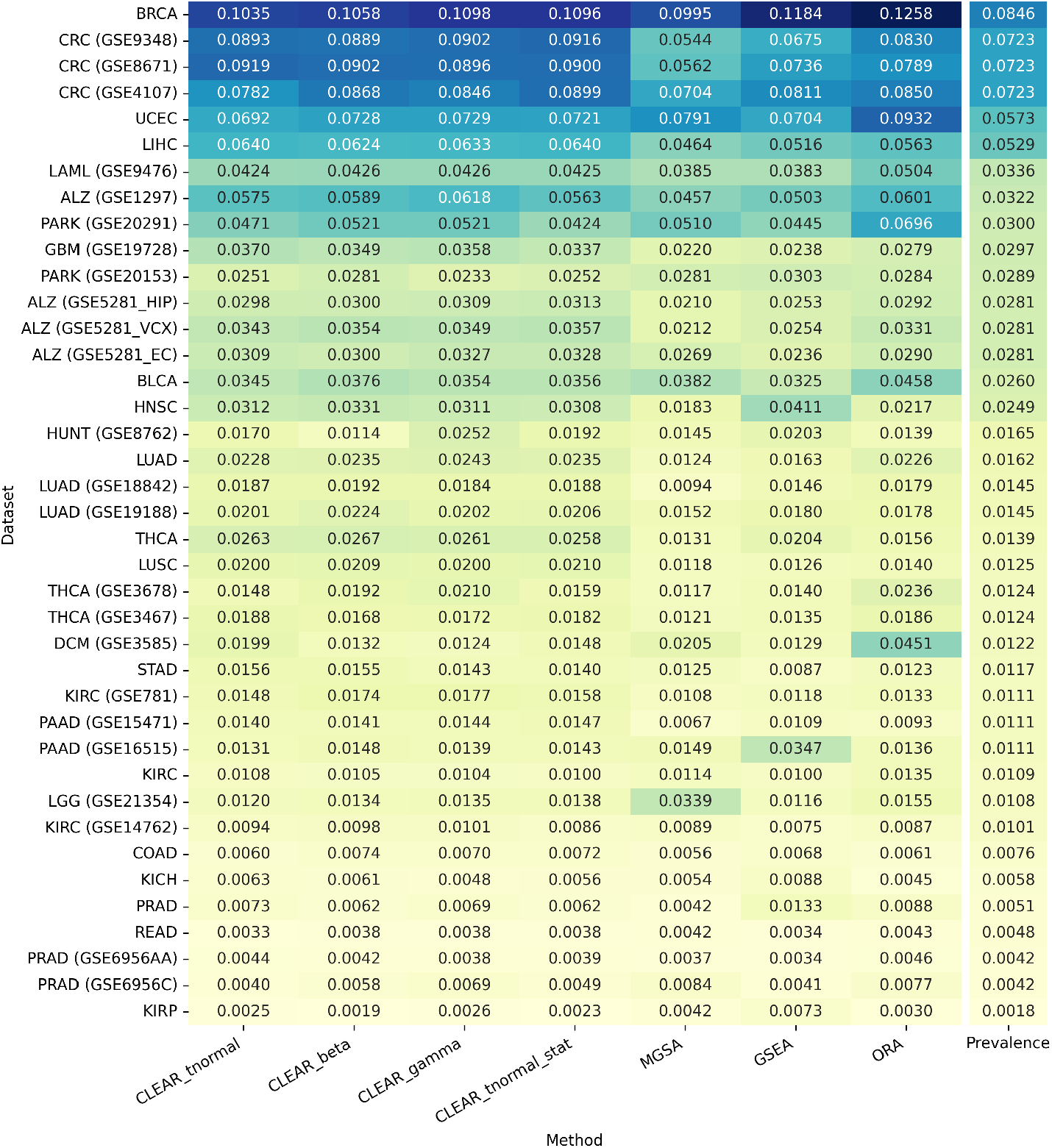
Heatmap showing area under precision-recall curve (PR-AUC) for all methods across 15 RNA-seq datasets (from TCGA) and 24 microarray datasets (from GEO, dataset ID is displayed), along with the prevalence of each dataset. Datasets are sorted by highest prevalence value. More than one dataset can be associated with the same type of cancer, and each cancer type has a unique set of phenotypically relevant gene sets (prevalence values could still differ due to difference in the number of valid gene sets in each dataset).

**Figure S6:**
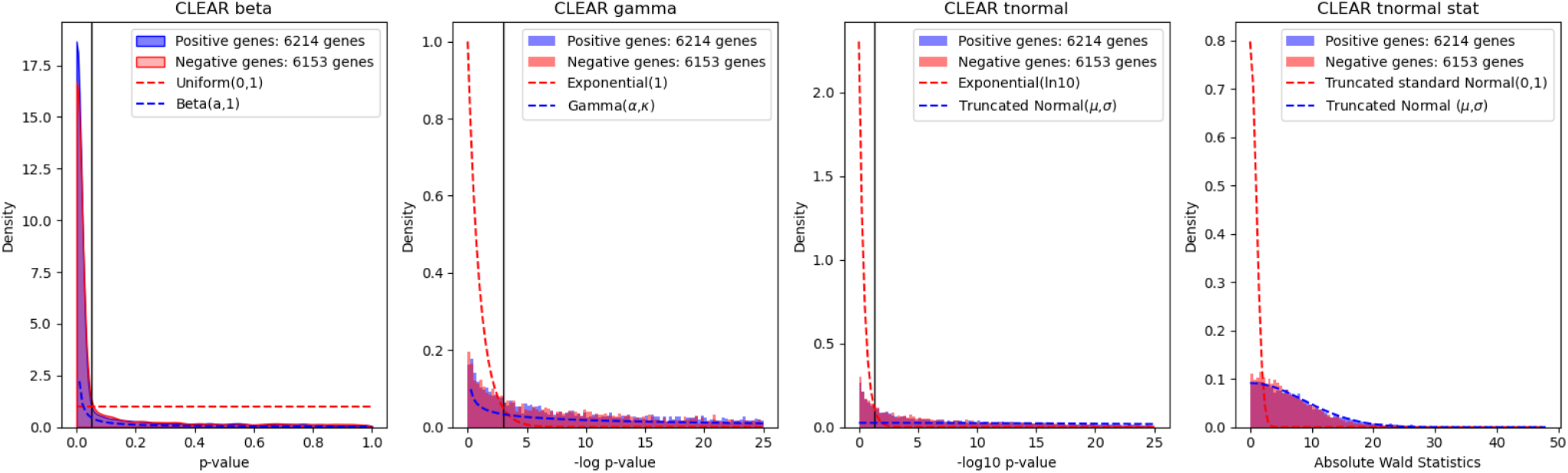
Gene-level statistic (*p*-value and Wald statistics) distributions of positive genes (genes belonging to cancer-relevant gene sets) and negative genes (remaining genes) in the BRCA dataset. Dashed red curves show the distributions under CLEAR’s null hypothesis assumption. Dashed blue curves show a fitted distribution for the alternative hypothesis, where parameters were set to their posterior means.

**Figure S7:**
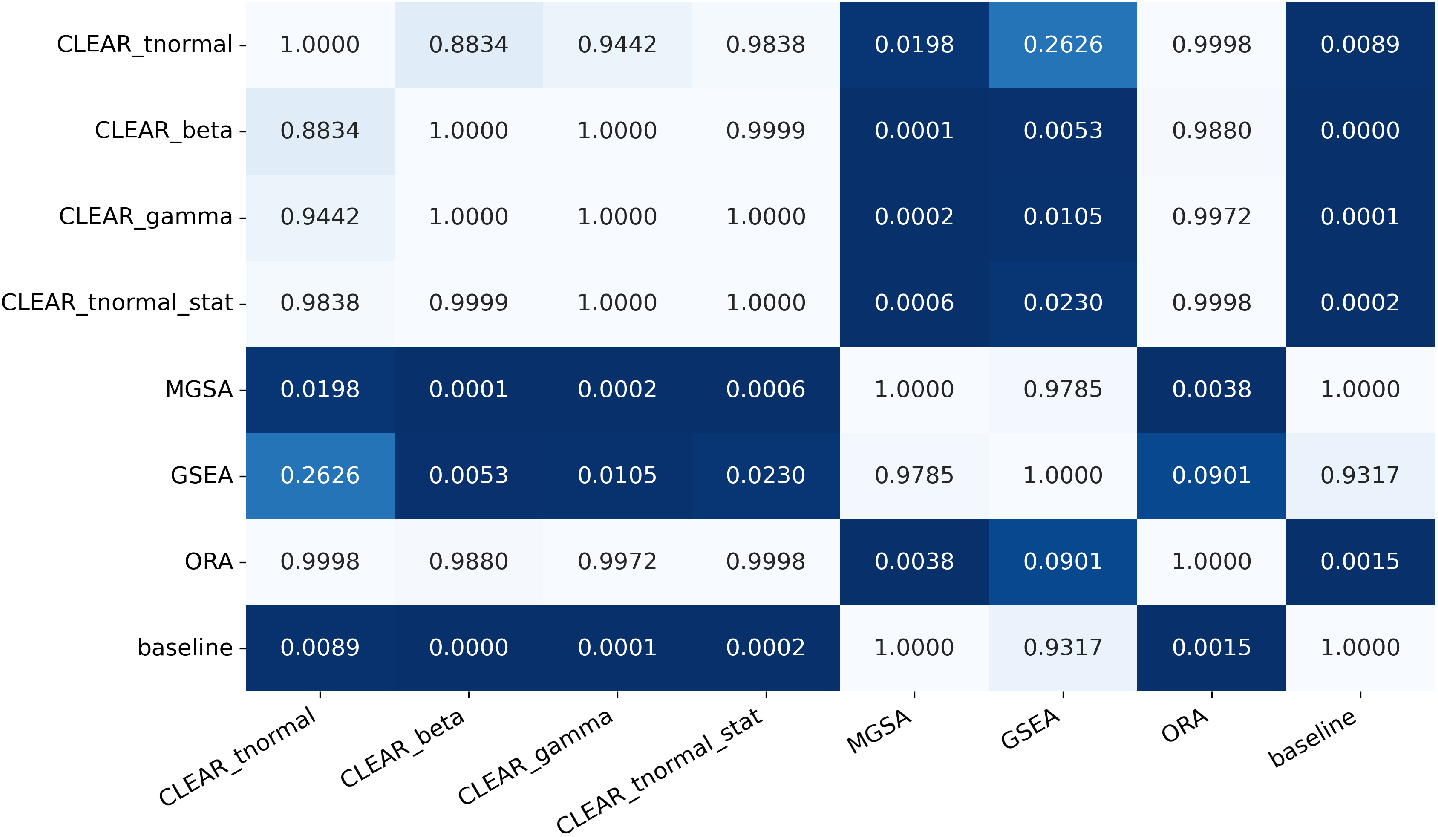
Pairwise post-hoc Nemenyi test *p*-values between normalized PR-AUC values of all methods. Baseline normalized PR-AUC value for each dataset is 1.0, which represents no gene set enrichment.

**Table S1:**
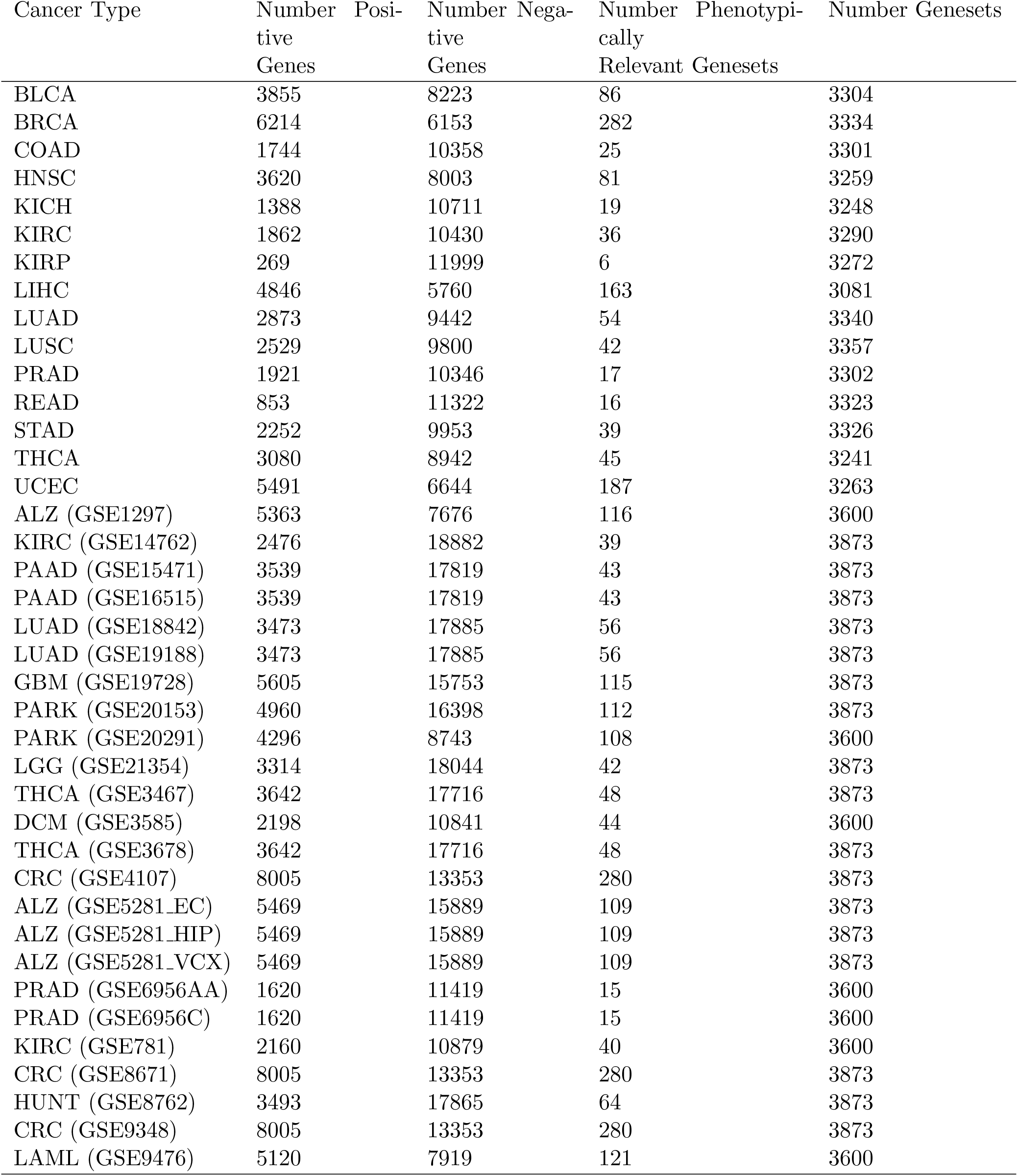
List of 15 TCGA RNA-seq datasets and 24 GEO microarray datasets. Phenotypically relevant gene sets were obtained from GSEABenchmarkeR package and were used to determine positive genes and negative genes. Num Genesets column shows the number of gene sets that have between 20 and 500 genes for each dataset.

